# The role of attention in the generation of anticipatory potentials to affective stimuli: an ERP and source analysis study

**DOI:** 10.1101/2025.07.10.664111

**Authors:** Ester Benzaquén, Timothy D. Griffiths, Sukhbinder Kumar

**Affiliations:** Biosciences Institute, Newcastle University, Newcastle Upon Tyne, United Kingdom

**Keywords:** anticipation, CNV, EEG, emotion, SPN

## Abstract

Anticipatory EEG signals are characterised by the occurrence of negative slow cortical potentials. This negativity is posed to be enhanced when expecting highly emotional stimuli; however, the specific role attention plays in its generation is unclear, as emotional content is more salient and arousing, and thus recruits higher attentional resources. Here, affective anticipation signals were recorded in 35 participants with EEG, and their brain sources elucidated using multiple sparse priors algorithm. Using a cued-paradigm, the category of a sound being negatively valenced or neutral could be predicted with a 68% accuracy. To shift attentional resources away from the emotional content, participants were instructed to listen and respond to a burst of white noise that occurred on 9% of trials. Results showed slower reaction times following the aversive cue, yet no difference in EEG amplitude between aversive and neutral anticipation. Response times positively correlated with EEG amplitude – participants with stronger negativity were faster to respond. EEG source reconstruction demonstrated no differences between conditions, and showed activation of areas within the salience network including insula, somatosensory cortex, and thalamus. The current results suggest that anticipatory EEG negativity is an index of attentional resource-allocation during the anticipation period and does not reflect the emotional content of upcoming stimuli.

---

The study of affect expectation has been dominated by fMRI (functional magnetic resonance imaging) research (e.g. see Ran et al. 2018), but due to its higher temporal resolution, EEG is more suited to measure the temporal dynamics of anticipatory activity. Anticipatory EEG signals are characterised by the occurrence of slow cortical potentials (SCPs) (see Birbaumer et al. 1990 for a review), generally of negative amplitude, that increase in magnitude as the anticipated event approaches. How emotions modify these SCPs appears to be an understudied and contentious area of research. Affective anticipation has commonly been studied using a first stimulus (S1) that signals the start of a fixed time interval until the presentation of a second stimulus (S2) of an emotional nature. Generally, the ERP component measured before S2 onset is called the Contingent Negative Variation (CNV) or the Stimulus Preceding negativity (SPN), depending on whether the S2 warrants a motor response or not, respectively. Anticipatory negative SCPs have been recorded before the presentation of affective stimuli, including threat stimuli such as shock (Böcker et al. 2001; Baas et al. 2002; Seidel et al. 2015; Tanovic et al. 2018), and affective images with negative (Takeuchi et al. 2005; Johnen and Harrison 2020) and positive (Simons et al. 1982; Poli et al. 2007) valence. However, previous research has found conflicting results, some showing an enhancement of negative SCPs before the presentation of emotional stimuli independent of valence (Qiao et al. 2018; Wiese et al. 2023), while others fail to replicate this (Takeuchi et al. 2005; Poli et al. 2007).

One problem with trying to measure and isolate affective anticipation by using emotional stimuli is that attentional processes are an inherent confound, as emotional content is typically more salient and arousing, and thus recruits higher attentional resources. This may be behind the lack of consistency in the literature thus far. For example, Takeuchi et al. (2005), using positive, negative, and neutral images, found an enhancement of anticipatory potentials before affective images of aversive valence but not for those of a positive valence. This was more than likely driven by differences in arousal, as aversive pictures were rated as significantly more arousing than pleasant ones. Poli *et al*. (2007) manipulated both valence and arousal and found enhanced anticipatory signals for high arousal images regardless of valence.

Older studies already provided evidence of the modulatory role of salience and attention to affective anticipatory SCPs. Simons et al. (1979) recorded anticipatory EEG signals before the presentation of high arousal nude images. Importantly, this anticipatory negativity was enhanced when the images were presented briefly (500 ms), and thus under higher attentional demand. When the images were presented for 6 seconds, the previously measured anticipatory SCP was blunted and did not show the traditional ‘ramping-up’ over time – in this condition, anticipatory attention is less essential, as the presentation of the stimulus serves as a bottom-up signal of when attentional resources can start to be recruited.

Attention is inherently intertwined with other preparatory processes and its specific effects are difficult to dissociate, even more so when using highly salient stimuli such as in affective research. Anticipation of a stimulus is likely to reflect both simple prediction of the upcoming stimulus/sensory input and neural adjustments to optimise behaviour by strategically focusing attention on task-relevant features. Attention is recognised to enhance sensory processing at the neural level, and this can occur well before stimulus onset (Driver and Frith 2000). It is also known, unsurprisingly, that a cue predicting a target will decrease response times (e.g. Fan et al. 2007). Associations between CNV amplitude and behavioural performance have been found numerous times (Haagh and Brunia 1985; Birbaumer et al. 1990; Wascher et al. 1996; Fan et al. 2007; Schevernels et al. 2014; Fishman et al. 2021), supporting the idea that task-preparation and the concomitant attention-allocation are indexed by the CNV, and thus an increased negativity signifies higher recruitment of attentional resources.

Another confounding factor which can bias the focus of attention in anticipatory SCPs relates to motivation of the subject to engage (or not) with the stimuli/task. For example, the SPN before performance feedback is significantly enhanced when a financial reward or punishment is added (Kotani et al. 2001; Kotani et al. 2003; Masaki et al. 2006; Ohgami et al. 2006; Schevernels et al. 2014). While some studies have managed to show a SPN while passively waiting for a monetary reward or punishment unrelated to performance (Donkers and van Boxtel 2005), others have shown the addition of an arbitrary outcome can blunt the SPN and almost entirely abolish it (Masaki et al. 2010). These seemingly opposite results may be explained if we consider participants’ motivational state. Attention will be high if only an arbitrary-outcome condition is included (Donkers and van Boxtel 2005). However, if besides this arbitrary condition participants are presented with an outcome contingent on participants’ performance, this would lower the motivational relevance of the arbitrary-reward condition by comparison (Masaki et al. 2010). These results highlight that the attentional state of participants plays an important role in SPN generation.

Even in the absence of a task, attention can be *re*oriented (or selective attention can be deployed) and prepared for processing of stimulus characteristics. It is likely that anticipatory SCPs reflect the specific characteristics of the stimulus that is about to be processed. For example, the SPN can have different scalp topographies, and thus different sources, depending on the nature of the anticipated stimulus (Ohgami et al. 2014). Exploring the neural sources of these SCPs can further delineate the mechanisms involved in their generation. Early attempts to localise EEG sources used dipole fitting models, which may lack in accuracy. A spatiotemporal dipole model implicated the insular cortex in the generation of the pre-feedback SPN (Böcker et al. 1994). The same group (Böcker et al. 2001), using a single dipole model, localised a frontocentral cortical negativity during the threat of shock to the ACC (anterior cingulate cortex).

Neural generators of slow EEG waves can also be studied using neuroimaging techniques such as fMRI or PET (positron emission tomography). Several studies appear to indicate that negative SCPs resemble the activity measured by increases of the hemodynamic response (reviewed by Khader et al. 2008). In a PET study comparing (knowingly) false with real performance feedback, Brunia and colleagues (2000) found right activation of the inferior frontal gyrus, posterior insula and inferior parietal lobe. Further, they found learning effects (increased activation in late vs early trials) in SMA. Tsukamoto et al. (2006), using a similar paradigm in fMRI, found activation of anterior and posterior insula, middle frontal gyrus, thalamus, and striatum during informative (i.e., true) feedback. Kotani and colleagues (2009) implicated again the insular cortex during feedback anticipation. Additionally, activation of the right anterior insula increased parallel to task difficulty. In a later study, the authors used the same paradigm in two different subsets of participants undergoing either EEG or fMRI measurements and manipulated whether performance feedback was presented (Kotani et al. 2015). Participants rated the feedback condition as more positively emotional and higher in arousal. fMRI activation was found in areas including ACC, occipital gyrus, MFG (middle frontal gyrus), ACC and MCC (mid-cingulate cortex), thalamus, right middle temporal gyrus, SMA, and insula. Subjective emotional scores correlated with activity in right anterior insula, right MFG, and right inferior parietal lobule. An fMRI-constraint source analysis of the SPN differences between the feedback and no-feedback conditions found dipoles originating in the anterior insula, ACC, MCC, and right MFG among others.

Here, a simple S1-S2 paradigm where cues (S1) predict with 75% validity the category of the upcoming sound (S2), which is either neutral or aversive, is used. Unlike previous studies that suffer from low number of participants – for example, Takeuchi et al. (2005) tested 11 participants, while Donkers and van Boxtel (2005) recruited 16 participants – we recorded anticipatory signals in a sample of over 30 people. Previous studies failed to account for the role of attention in the generation of affective anticipatory potentials. Here, attention is controlled for by the introduction of catch trials that require a motor response, which happen equally in the affective and neutral condition. Although arousal is inherently different between aversive and neutral sounds, this is minimised by directing participants’ attention to an irrelevant task unrelated to the affective valence of the stimuli. To localise the sources of the anticipatory potentials, a Bayesian approach is used – Multiple Sparse Priors (Friston et al. 2008) – which is known to perform better than previous models (Hyder et al. 2014; Jatoi et al. 2016). If anticipatory potentials do not solely represent recruitment of attentional resources, but are also an index of emotional anticipation, recorded cortical potentials will differ in amplitude and/or sources according to the valence of the predicted stimuli. On the other hand, if attention is the main or sole determinant of these SCPs (SPN/CNV), amplitude will be modulated by successful recruitment of attentional resources as measured by reaction times to catch trials.

## Materials and Methods

### Participants

Data from 35 participants (14 males) were analysed. Participants’ age ranged from 18 to 40 years (mean ± standard deviation [SD]: 25.67 ± 7.16). To take part, participants were required to be right-handed, aged between 18-40, have normal hearing and normal or corrected-to-normal vision, no history of neurological, psychological, or psychiatric disorders, and to not be taking any psychotropic drugs or neuroactive medication. The study was approved by Newcastle University’s Faculty of Medical Sciences Ethics Committee (Reference number: 1418/732/2017), and written informed consent was obtained from all participants before the start of the study. All volunteers were provided with an inconvenience allowance of £15 for their time.

### Experimental paradigm

Two visual cues preceded the presentation of two 1-second-long sounds: either a ‘neutral’ water sound (brook) or an ‘aversive’ scratching sound (fork on bottle). The aversive sound was a scratching sound perceived as highly aversive of a fork scratching a bottle adapted from Kumar et al. (2008), while the neutral sound was a shortened version of sound 172 (Brook) from the IADS-2 (Bradley and Lang 2007), adapted from Kumar et al. (2008). Normative subjective ratings of these sounds (2-seconds long) from Kumar et al. (2008) are 7.84 and 0.85 for the scratching and water sounds, respectively, on an unpleasantness scale 0-9. In a separate unpublished study (n=22), in an unpleasantness scale 0-10, these adapted sounds (1-second long) were rated as 8.9 ± 1.75 and 0.85 ± 1.20, respectively. The cues were the letters ‘X’ or ‘Y’ and had a 68.2% predictive validity, meaning each cue was followed by one of the sounds 68.2% of the time (congruent trials), and followed by the other sound 22.7% of the time (incongruent trials). On the remaining trials (9.1%), the cues were followed by a ‘catch-sound’. The association of each cue with either the neutral (water) sound or the aversive (scratching) sound was counterbalanced across participants. Participants were informed one of the cues tended to be followed by the aversive sound, but the exact probabilities and the identity of said cue were not provided. The cue was presented one second before sound onset and remained on screen while the sound was playing such that its offset aligned to sound offset. A new trial started between 1.3 and 1.5 seconds after sound offset. The trial structure is depicted in Figure 1.

**Figure 1.**
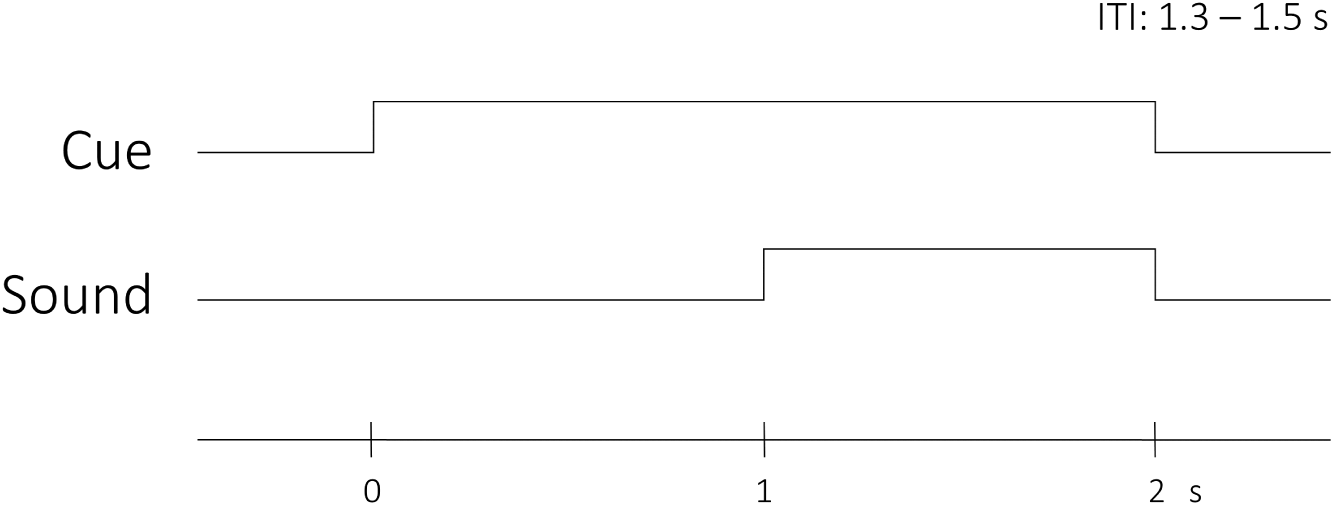
Trial structure. The visual cue was presented one second before sound onset for a total of two seconds. Sound and cue offset were aligned, and a new trial started 1.3 – 1.5 seconds after offset (ITI: inter-trial interval).

Cues were presented in equal proportions in a pseudorandomised order with no more than 6 sequential repetitions for a total of 600 experimental (neutral or aversive) trials. Thus, from 300 trials of the aversive cue, the aversive sound was presented on 225 trials, while the neutral sound was presented for a total of 75 trials in a randomised order with several constraints; mainly, no incongruent trial appeared in the first 3 presentations of each cue, and no more than 3 sequential incongruent trials after each cue were allowed. An extra 60 catch trials to control attention were added where the aversive or neutral sound was replaced by white noise (30 per cue), and participants were instructed to make a response by pressing the space bar as quickly as possible within one second. After a response, performance feedback (reaction time) was shown on screen for 1.5 seconds. If participants failed to respond or had a reaction time slower than 500 ms, the message ‘Please respond faster’ appeared on screen. After few response trials, the RT threshold for this message changed from 500 ms to the mean RT + 2 standard deviations. To proportionally distribute catch trials along the experiment, 2 catch trials were randomly added for each 20 experimental trials. Participants practiced the task with a shortened version of only 54 trials. The task was divided into four blocks and time of rest was offered between blocks. At the end of the task, participants were asked which sound tended to follow each cue to ascertain the cue-sound association was indeed learnt.

### Behavioural analysis

All reaction times to catch sounds (i.e., white noise) were standardized within each participant. These were then averaged according to whether the preceding cue predicted ‘mostly’ aversive or neutral sounds, and a paired two tailed t-test was performed.

### EEG preprocessing

EEG was recorded using a 64-channel ActiveTwo BioSemi system (BioSemi Instruments, Amsterdam, The Netherlands) at a sampling rate of 1024 Hz. Vertical and horizontal EOG (electro-oculogram) signals were recorded from electrodes positioned above and below the right eye and on the outer canthi of both eyes. Electrode DC offset (akin to impedance) was kept stable and within ±30 mV. Data were epoched to cue onset from −1.29 to 3 seconds, and robust detrended (de Cheveigne and Arzounian 2018) to remove slow drifts instead of the standard application of a highpass filter, as filtering may introduce distortions in the data (Tanner et al. 2015; Yael et al. 2018). Ocular-artifacts were identified using AMICA (Adaptive Mixture Independent Component Analysis) (Palmer et al. 2011), which achieves a better ICA decomposition compared to other algorithms such as Infomax (Delorme et al. 2012). After rejecting EOG artifacts, EOG channels were removed and common average referencing was performed. Baseline correction from 200 ms before cue onset was applied. Data were then converted into Fieldtrip format, and lowpass filtered to 30 Hz.

To study anticipatory potentials (SPN/CNV), data were divided according to visual cues into aversive and neutral trials, and trial rejection was performed for each condition separately using Fieldtrip’s visual rejection tool and computing variance. Epochs were time-locked to the cue – from 0.1 before to 1 second after cue onset. Trials with gross artifacts were visually identified as outliers and removed. When noisy channels were detected this way, they were also removed and interpolated using ‘neighbours’ created with Fieldtrip’s ‘biosemi64’ template with the weighted method. If 30% or more of the trials were removed this way, for any of the two conditions, the participant was rejected and excluded from any EEG analysis. A second trial rejection was further performed where trials with a high variance (> 130) were automatically rejected. This criteria was visually selected as successful in the removal of skin potentials. Due to a great number of skin potentials and to retain as much data as possible, this trial rejection was performed for each channel separately, meaning a noisy trial caused by artifacts on a small selection of channels did not need to be rejected in all of the channels. For this reason, rejected trials for each channel were replaced by NaNs to maintain the same trial structure for all channels. If less than 200 trials remained after trial rejection on a given channel per condition (i.e., cue), this channel was removed and interpolated. If more than 10 channels were interpolated for a given participant, this participant was excluded from further EEG analyses (but retained for solely behavioural [RT] analysis). Using these criteria, a total of 4 participants were rejected due to artifacts and an average of 0.87 ± 1.62 and 0.84 ± 1.53 (mean ± SD) channels were interpolated for the aversive and neutral conditions respectively. During the first trial rejection, an average of 8.81 ± 5.31 and 7.68 ± 5.50 % of trials were rejected for the aversive and neutral conditions respectively. During the second trial rejection, a further 1.12 ± 1.22 and 1.20 ± 1.29 % of trials were removed. In total, an average of 9.92 ± 5.92% and 8.88 ± 6.22% of all aversive and neutral trials were rejected.

### EEG analysis

#### Anticipatory potentials (SPN/CNV)

Anticipatory potentials, termed SPN from now on for ease of understanding, were calculated by averaging the EEG activity over the last 200 ms before sound onset, consistent with previous research (e.g., Brunia and Damen 1988; Poli et al. 2007; Tanovic and Joormann 2019). Differences between conditions over all channels were assessed using a Monte Carlo permutation test on Fieldtrip with 1000 permutations and a (spatial) cluster correction with a significance threshold of *p* < .05. To further characterize the SPN, a single-channel analysis was performed. To determine the most responsive channel to the expectation of upcoming sound without bias, the SPN was averaged between all conditions and participants, and the channel with the minimum amplitude was selected for further analysis. Differences between conditions on this channel were tested using Fieldtrip’s dependent-sample permutation test with *p* < .05.

#### Relation between SPN and behaviour

The SPN (i.e., the last 200 ms before sound onset) was averaged over time and between conditions (i.e., cues) for each participant. The channel with the strongest negativity after averaging the SPN among participants was selected for analysis. Pearson correlation was performed between the mean amplitude of the SPN for all trials on the selected channel and the averaged reaction times to catch sounds regardless of cue. Two further correlations were performed by splitting the SPN and RTs depending on the category of the cue to explore whether the relationship between RTs and anticipatory potentials was modulated by emotion.

#### SPN source reconstruction

Source analysis of the anticipatory activity was performed in SPM12 (https://www.fil.ion.ucl.ac.uk/spm/software/spm12/). The SPN during the last 200 ms before sound onset was averaged over trials for each condition and subject. SPM’s head model based template in the MNI space was used for co-registration. The EEG-BEM (Boundary Element Method) forward head-model was selected to compute the lead field matrix. The forward model was inverted using the Multiple Sparse Priors (MSP) algorithm using greedy search (Friston et al. 2008). MSP is a Bayesian approach to the inverse problem which automatically selects multiple cortical sources using empirical priors: it calculates covariance components of each prior and optimizes a model using restricted maximum likelihood. A group inversion was performed to restrict the activated sources between participants (Litvak and Friston 2008). A one-Sample t-test using SPM statistics with a *p* < .001 (FDR cluster-corrected) was used to define statistically significant sources of activation per condition. A paired t-test was performed to identify differences in sources between conditions. Anatomical labels were extracted using the automated anatomical labelling atlas 3 (AAL3) (Rolls et al. 2020).

## Results

### Behavioural

All participants correctly identified which sound tended to follow each cue when asked after the experiment. No outlier reaction times (<150 ms or >1) were observed, and thus, all RTs were included in the analysis. Normalized reaction times to the white noise (catch trials) after the aversive cue were significantly slower than responses after the neutral cue (aversive: 418.84 ± 59.78 ms; eutral: 408.21 ± 57.41; *t*(34) = 2.68, *p* = .011; Figure 2).

**Figure 2.**
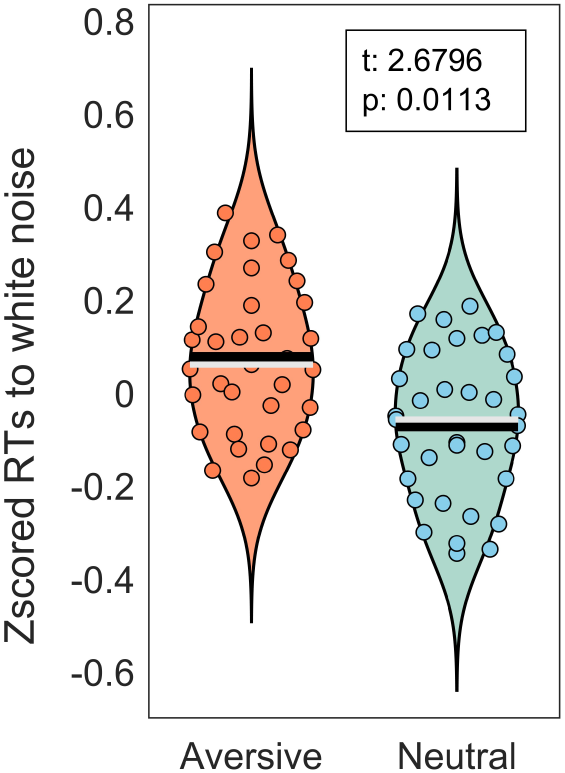
Normalized response times to the white noise after aversive and neutral cues. Black and white lines indicate the mean and median respectively.

### Anticipatory signals (SPN/CNV)

SPN did not differ between cues; that is, the amplitude of the SPN was similar regardless of the cue in all channels (all *ps* > .05, cluster corrected). The scalp topography of the SPN before sound onset can be seen in Figure 3A. The SPN over the channel (Cz) which showed maximal anticipatory amplitude also did not differ between conditions (mean ± SD; aversive: −1.31 ± 1.64; neutral: −1.26 ± 1.79; *p* = .22; Figure 3B and Figure 3C).

**Figure 3.**
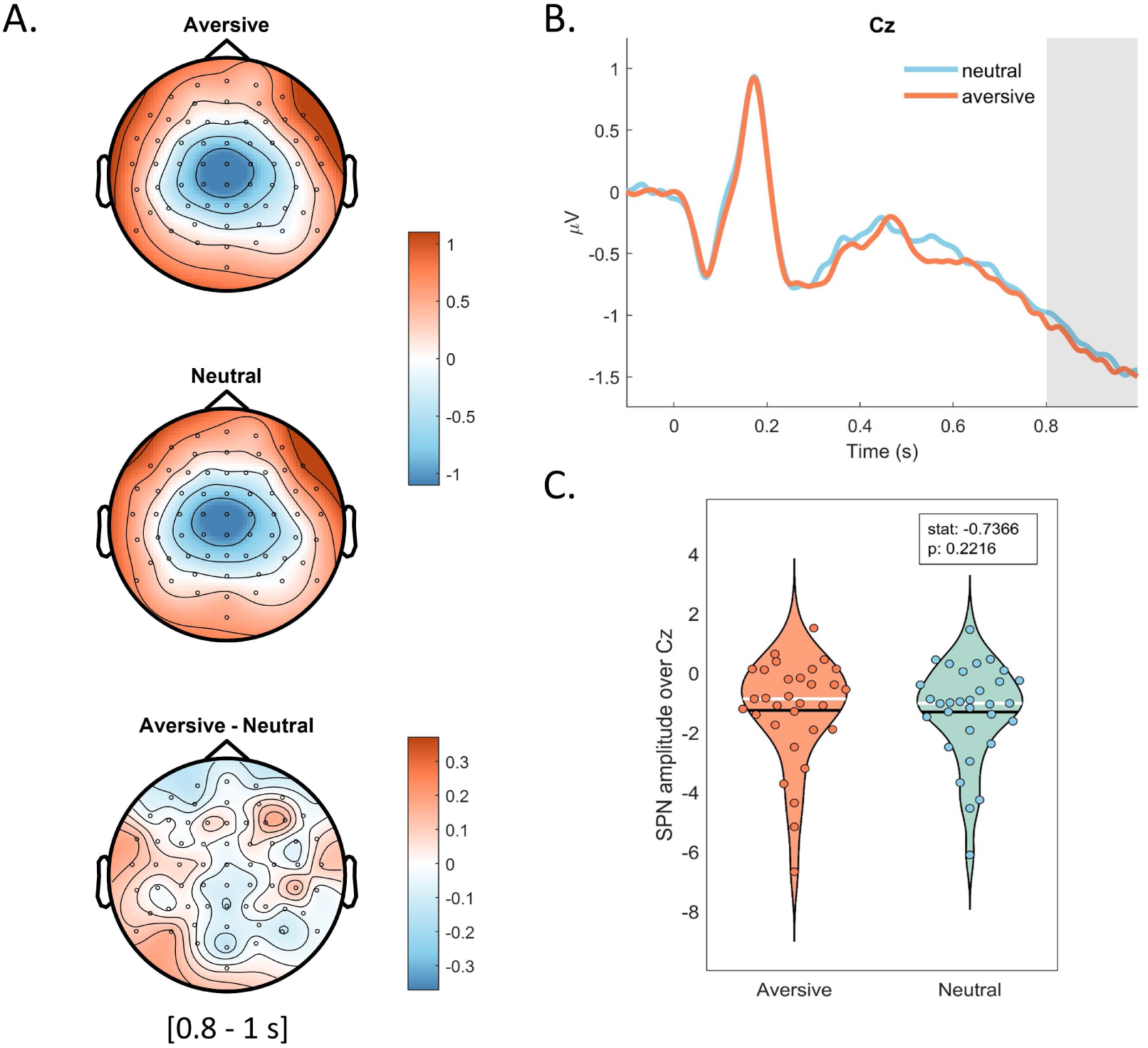
SPN differences between conditions. A: Topographical maps of the anticipatory period [200 ms before sound onset] after aversive and neutral cues and their difference (all p > .05, cluster corrected). B: Cue-locked ERPs on channel Cz; studied time window highlighted in grey [last 200 ms]. C: Averaged SPN amplitude over maximally negative electrode (Cz) for each participant and condition (p > .05, permutation test). White and black lines represent the median and mean, respectively.

### Relation between behaviour and SPN

A significant positive correlation was found between participants’ averaged reaction times to catch trials and SPN amplitude on channel Cz (Pearson’s r: .69, *p* < .001, n = 31; Figure 4). Participants who exhibited a stronger SPN overall had faster response times during catch trials. This relationship remained after splitting trials based on cues (Neutral condition; r: .69, *p* < .001; Aversive condition; r: .67, *p* < .001).

**Figure 4.**
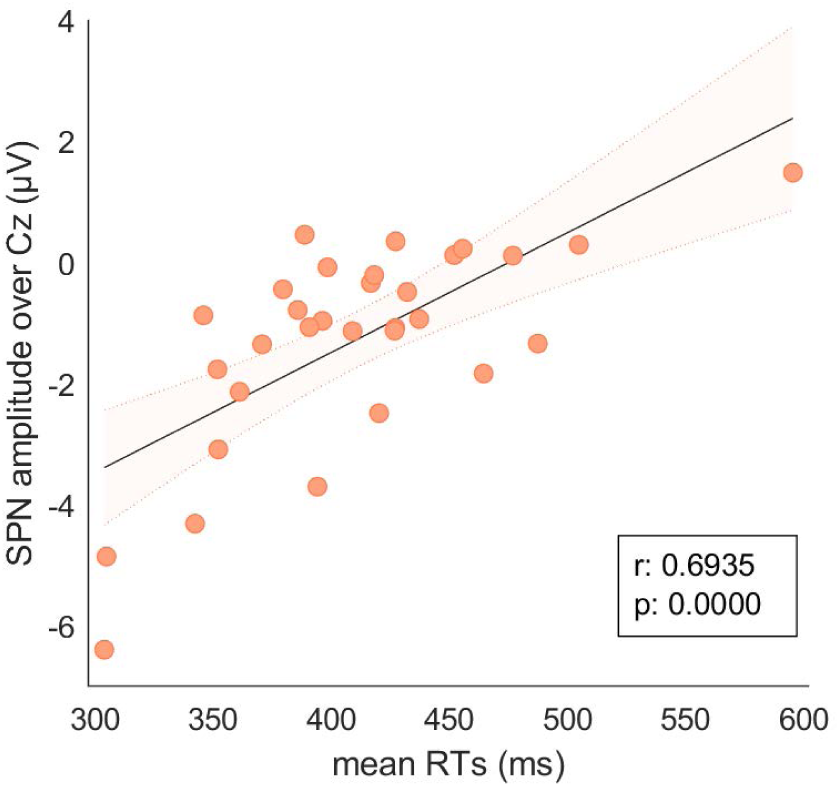
Pearson’s correlation between RTs in ms and anticipatory (SPN) amplitude on Cz.

### SPN source reconstruction

Source reconstruction did not differ between conditions (cue type). One-sample t-tests results for aversive and neutral trials displayed identical source solutions, thus only aversive results are reported. Source analysis revealed anticipatory activation of frontal, temporal, somatosensory and visual cortices. Areas included bilateral anterior insula, IFG (inferior frontal gyrus), right MFG (middle frontal gyrus), SMA, parts of the thalamus and right ACC, and superior, middle and inferior temporal gyri. Source space activation can be seen in Figure 5. A full list of the main areas in each cluster can be found on Table 1.

**Figure 5.**
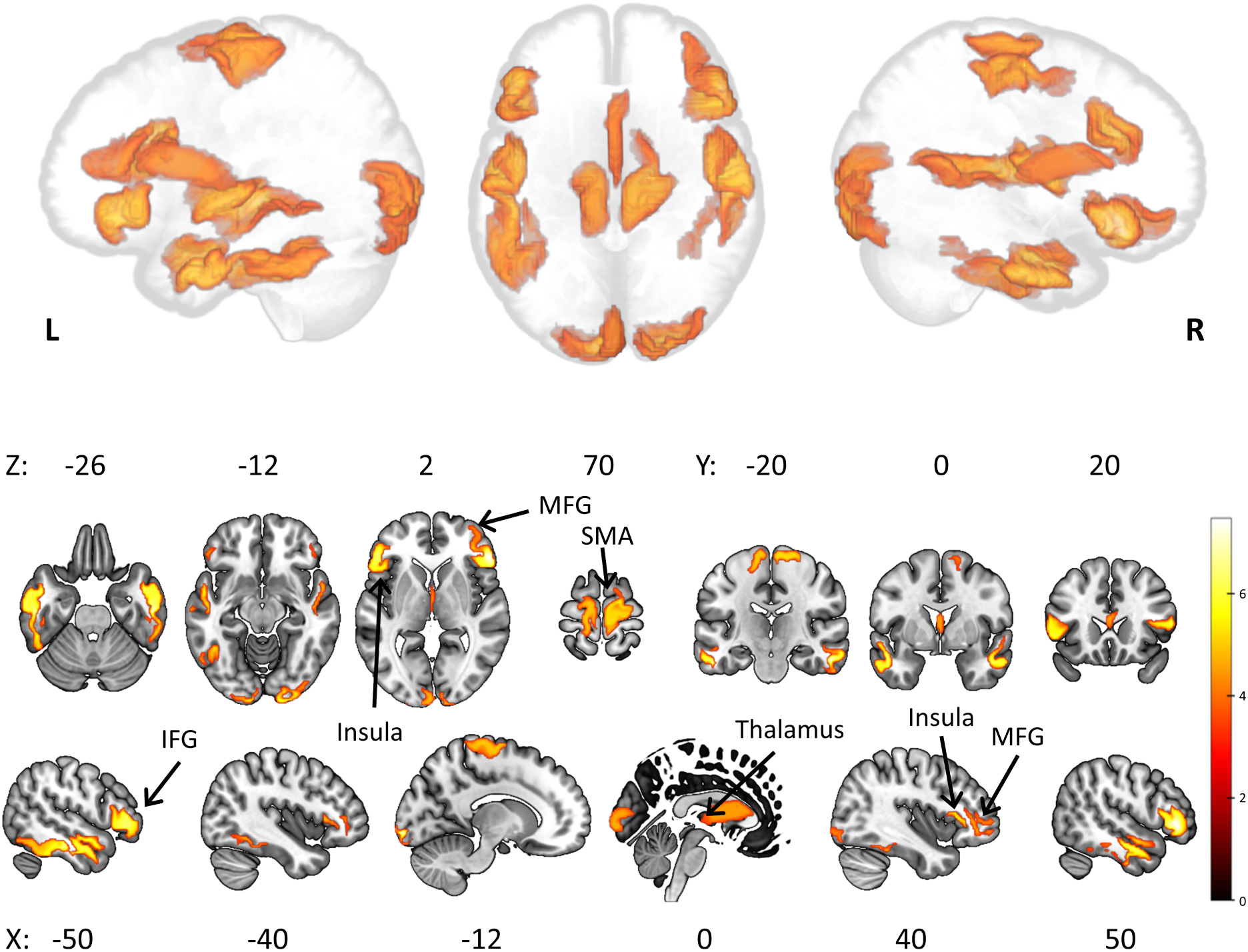
Source reconstruction results projected on glass brain (top) and MNI template (bottom). SPN anticipatory activity was localised to temporal and occipital cortices, anterior insula, IFG, SMA, ACC and midbrain. L: left; R: right.

**Table 1.**
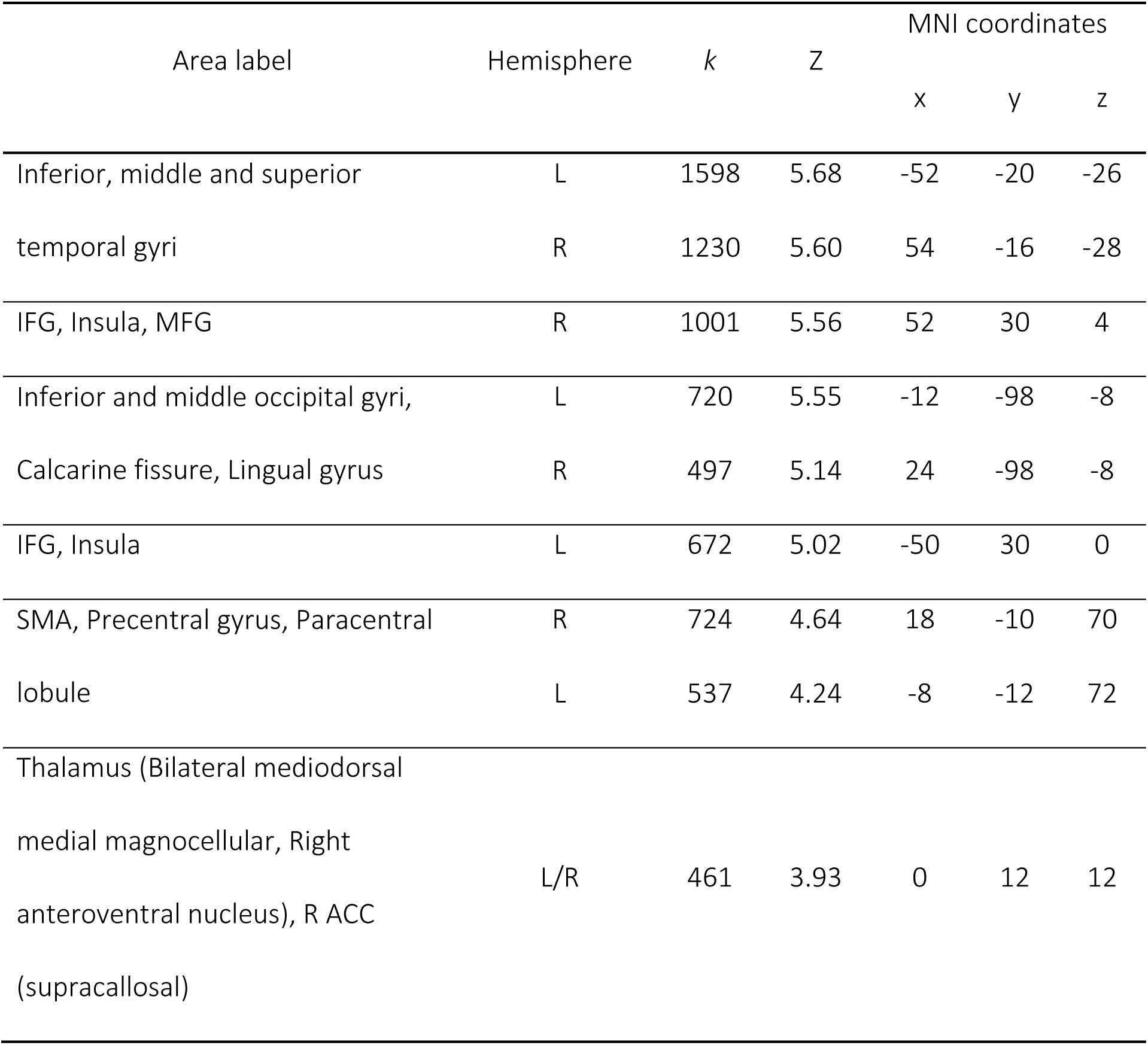
Cluster results of SPN source analysis.

## Discussion

In the current study, anticipatory signals to affective sounds were recorded with EEG. Using a cued-paradigm, the category (valence) of sounds could be predicted with a 68% accuracy. Participants were instructed to listen and respond to a white noise that occurred on 9% of trials (i.e., catch trials). The ‘SPN’ [last 200 ms before sound onset] was explored during aversive and neutral anticipation, and its sources assessed using multiple sparse priors algorithm (Friston *et* al. 2008).

Reaction times to catch trials were slower after an aversive cue compared to when the cue predicted neutral sounds. Unlike avoidable threat, which elicits motor preparation, inescapable aversive events are characterized by “attentive freezing”, which is accompanied by potentiation of skin conductivity and the startle reflex, and fear bradycardia (Löw et al. 2015), which could explain the slowing of reaction times in our data. However, we found no differences in the EEG amplitude during the anticipation of aversive or neutral sounds. Further, response times to catch trials positively correlated with EEG amplitude – participants with overall stronger (i.e., more negative) anticipatory negativity were faster to respond. The relation between RTs and ‘SPN’ amplitude underlines the importance of selective attention or attentional vigilance to the generation of anticipatory potentials. If participants’ attentional resources were directed towards the target sound, and the recorded pre-stimulus negativity does not reflect emotional anticipation but only attention/motivation, the anticipatory activity would indeed not be modified by the cue. It could be argued that there was no emotional expectation induced - that participants’ anticipatory activity was only related to the ‘catch’ white noise and not the emotional stimuli. For example, actively predicting the catch sound regardless of cue might pose a strategic advantage compared to anticipating aversive or neutral sounds, especially considering cue contingencies were not completely accurate – cues were only predictive of the sound on 75% of trials. Notwithstanding, the RT difference between cue types suggests participants anticipated aversive sounds after the aversive cue on a trial-by-trial basis, but this did not affect anticipatory potentials. Further, post-experiment questions revealed participants learnt the cue-sound contingencies. Thus, taken together, it is likely that affective expectations were induced. The current results indicate that the pre-stimulus negativity (SPN/CNV) is an index of motivational anticipation ruled by attentional resource-allocation and does not reflect the emotional content of upcoming stimuli.

Some research has found personal differences in the amplitude of affective anticipatory potentials, including high worry (Grant et al. 2015), intolerance of uncertainty (Gole et al. 2012; Wiese et al. 2023), autism spectrum disorder (Stavropoulos and Carver 2014), schizophrenia (Wynn et al. 2010), Parkinson’s disease (Mattox et al. 2006), and depression (Umemoto and Holroyd 2017). Under this view, attention-allocation and cognitive control might be behind these differences, and not emotional processing mechanisms.

Source reconstruction of anticipatory potentials did not differ between neutral and aversive conditions. Anticipation after the aversive cue showed activation of areas within the salience network, somatomotor, visual and temporal cortices, as well as the midbrain. The involvement of SMA and other somatomotor regions, insula, cingulate, and thalamus found in this source reconstruction is extensively supported by the literature on negative SCPs. Nagai et al. (2004) performed an fMRI study on preparatory motor responses using an S1-S2 auditory paradigm, and found activation of cingulate and somatomotor cortices, bilateral insula, left caudate and right DLPFC (dorsolateral prefrontal cortex) and OFC (orbitofrontal cortex). In a subset of participants, they recorded simultaneous EEG and found that the trial-by-trial fMRI activity was modulated by CNV amplitude in the thalamus, anterior cingulate, SMA, and the hindbrain. Hinterberger et al. (2003), using neurofeedback, trained participants to generate sustained negativity at the vertex (Cz). During an fMRI session, they localised the self-induced negativity to activation in pre- and post-central gyri, SMA, insula, MFG, and DLPFC. In a similar study with simultaneous EEG and fMRI measurements (Hinterberger et al. 2005), they found that in the preparatory period before regulation of scalp potentials was required, a central negativity was measured using EEG and its amplitude covaried with the BOLD response in thalamus and SMA. Additionally, the self-induced negativity during the biofeedback session was associated with increased SMA and thalamus activity. Mento et al. (2013), during a passive task, and despite the lack of a motor component, localised the SPN to the SMA, highlighting the role of SMA in stimulus anticipation that goes beyond motor preparedness. Lastly, these areas have been extensively implicated in anticipatory affect (reward or loss anticipation). In a meta-analysis including 12 independent fMRI studies on reward processing, Knutson and Greer (2008) found that ‘gain anticipation’ was related to greater activation when compared to ‘gain outcome’ in the Nucleus Accumbens (NAcc), insula, caudate, MFG and thalamus. Another meta-analysis (Liu et al. 2011) on reward anticipation similarly involved areas such as the NAcc, insula, thalamus, brainstem, putamen, SMA, and ACC. Additionally, the anterior insula is not only an area of affective anticipation but has been implicated in numerous timing tasks involving temporal estimations (Kosillo and Smith 2010), which supports a broader role in stimulus anticipation.

Other research on motor-preparatory activity has also implicated frontal, occipital and temporal cortices. For example, Fan et al. (2007) performed the same paradigm in participants who underwent either EEG or fMRI. Preparatory activity (pre-response), as measured by fMRI, was found in a thalamo-cortico-striatal network which included the ACC, pre-SMA, MFG, SFG (superior frontal gyrus), anterior insula, superior occipital gyrus, the caudate, and midbrain. The ‘post-cue’ (S1) CNV amplitude, as measured by EEG and which correlated with faster response times, was localised using dipole modelling whose locations were constraint based on the fMRI results (excluding putamen and caudate), and could explain 66% of the ERP variance. The anterior insula and occipital cortex have also been found active during the anticipation of shock and aversive images (Simmons et al. 2004; Seidel et al. 2015). Although at first occipital activation during sound anticipation was unexpected, previous literature has involved the visual cortex in attentional shifts towards auditory stimuli (McDonald et al. 2013). Involvement of the visual cortex during anticipation of sounds was also found by Bueti and Macaluso (2010) during a simple motor task, as well as sensorimotor regions, insula, and thalamus. Prefrontal and motor cortices have been involved in forming temporal expectations (Coull 2009), and direct recordings from non-human primates show involvement of neurons in prefrontal and cingulate cortex during anticipation (Niki and Watanabe 1979). The motor cortex thus appears to be involved in the processing of timing of events or temporal learning which is further supported by the literature (Allman et al. 2014).

The engagement of temporal areas during anticipation in the current study, which is not found by previous affective research – which usually use visual stimuli – indicates this is most likely due to the use of auditory stimuli; thus, auditory cortices appear active before sound presentation. This may represent resource allocation to auditory cortices in preparation for sound processing (e.g., Gómez et al. 2004). Alternatively, this activation might be due to sustained attention to ‘internal representations’ of the target or upcoming sound, since the storing and retrieval of sounds in working memory have been shown to activate the auditory cortex (Huang et al. 2016). This is further supported by research showing increased EEG anticipatory negativity before predicted word and speech onset can be localised to the temporal cortex (Grisoni et al. 2024). Taken together, the current source reconstruction of anticipatory potentials includes areas that have been previously related to anticipation, and thus add evidence to the veracity of these results.

### Limitations

A major limitation of the current study is its inability to dissociate between SPN and CNV. Although SPN and CNV are highly related, and some researchers do not differentiate between them (e.g., Mento et al. 2013; Del Popolo Cristaldi et al. 2021), SPN is considered to reflect only stimulus expectancy, while CNV includes motor preparedness. By adding catch trials, and thus requiring a motor response, the measured anticipatory ERP could be classed as CNV. Nevertheless, the reported anticipatory ERP, whether CNV or SPN, includes stimulus expectancy, and this was not modulated by the emotional content of the upcoming stimuli. However, the fact that we cannot differentiate between SPN and CNV leaves out the possibility that the two processes, if they are indeed categorically different, might be affected by emotion and attention differently. Further work on solely anticipatory signals without a motor component would help answer this question. Additionally, the considerably low predictive value of each cue (68%) might have altered our findings. Anticipatory potentials can be modulated by uncertainty (Johnen and Harrison 2020), and thus it may be that the lack of a higher predictability induced a degree of uncertainty that abolished the otherwise measurable affective effect on anticipatory potentials. Lastly, due to the use of aversive stimuli of high arousal in the current study, any hypothetical modulation in anticipatory potentials could have been driven by changes in arousal and not emotional content. Future research where attention, arousal, and emotional content are orthogonally modified without necessitating a motor response at the time of stimulus presentation could further elucidate the role of attention in the generation of the affective SPN.

## CRediT author statement

Ester Benzaquén: Conceptualization, Data Curation, Formal analysis, Investigation, Methodology, Visualization, Writing - Original Draft.

Timothy D. Griffiths: Conceptualization, Resources, Writing - Review & Editing.

Sukhbinder Kumar: Conceptualization, Funding acquisition, Methodology, Writing - Review & Editing.

## Funding

This work was supported by Newcastle University. The sponsor had no involvement in the study design, collection, analysis and interpretation of data, writing of the report and decision to submit the article for publication.

## Data availability

The data underlying this article are available in OSF (Open Science Framework) at https://dx.doi.org/10.17605/OSF.IO/8PZVK.

## Notes

### Competing Interest Statement

The authors have declared no competing interest.

